# Variation in hemolymph content and properties among three Mediterranean bee species

**DOI:** 10.1101/2023.12.22.573070

**Authors:** Salma A. Elfar, Iman M. Bahgat, Mohamed A. Shebl, Mathieu Lihoreau, Mohamed M. Tawfik

## Abstract

Hemolymph, as mediator of immune responses and nutrient circulation, can be used as physiological marker of an insect’s health, environmental quality or ecological adaptations. Recent studies reported intraspecific variation in protein contents and biological activities of the hemolymph of honey bees related to their diet. Here we measured interspecific variation in three common bee species in the Mediterranean Basin with contrasting ecologies: *Apis mellifera*, *Chalicodoma siculum,* and *Xylocopa pubescens*. Despite all the bees were collected in the same area, we found important quantitative and qualitative variations of hemolymph extracts across species. Samples of *A. mellifera* and *C. siculum* had much higher protein concentration, anticancer, antimicrobial and antoxidant activities than samples of *X. pubescens.* This first descriptive study suggests life history traits of bee species have strong influences on their hemolymph properties and call for future large scale comparative analyses across more species and geographical areas.

## 1. Introduction

Bees are a large and diverse taxonomic group of about 20,000 species categorized in seven families [1]. Many of these species provide well-known natural products such as honey, bee pollen, wax, royal jelly and venom that have antioxidant, antibacterial or antitumor activities [2]. Increasing evidence suggest bee hemolymph also possesses interesting biological properties [3–6].

Hemolymph is a vital fluid involved in nutrient circulation to nourish tissues and in immune responses to fight infections [7]. Bee hemolymph contains various hydrophilic (e.g. hemocyanin) and hydrophobic (e.g. apolipophorins) proteins [8–10]. High concentrations of these proteins improve the immune responses of bees and their resistance to infections [11–13]. Recent work described significant intraspecific variations in the proteomic profiles of the hemolymph of bees that are likely related to the diet [6,14]. Therefore, it has been proposed that the proteomic structure of hemolymph can be used to monitor the physiological conditions of bees as well as the quality of their environment. Since bees are found across a wide range of habitats worldwide [1], we also expect hemolymph variation across species to exist and reflect key ecological adaptations.

Here we explored interspecific variations in hemolymph properties of bees from the Mediterranean Basin. We analysed hemolymph samples of three common species collected in the same area in Egypt: *Apis mellifera*, *Xylocopa pubescens* and *Chalicodoma siculum* (Figure 1). *A. mellifera* (honey bees) are social Apidae nesting in cavities [15]. *X. pubescens* (carpenter bees) are solitary Apidae that build their nests in wood [16,17]. *C. siculum* are solitary Megachilidae that build their nests with mud [18]. This is a descriptive study that primarily aimed at analyzing honey bee hemolymph and comparing it to that of two other common yet poorly studied bee species (*C. siculum* and *X. pubescens*).

**Figure 1.**
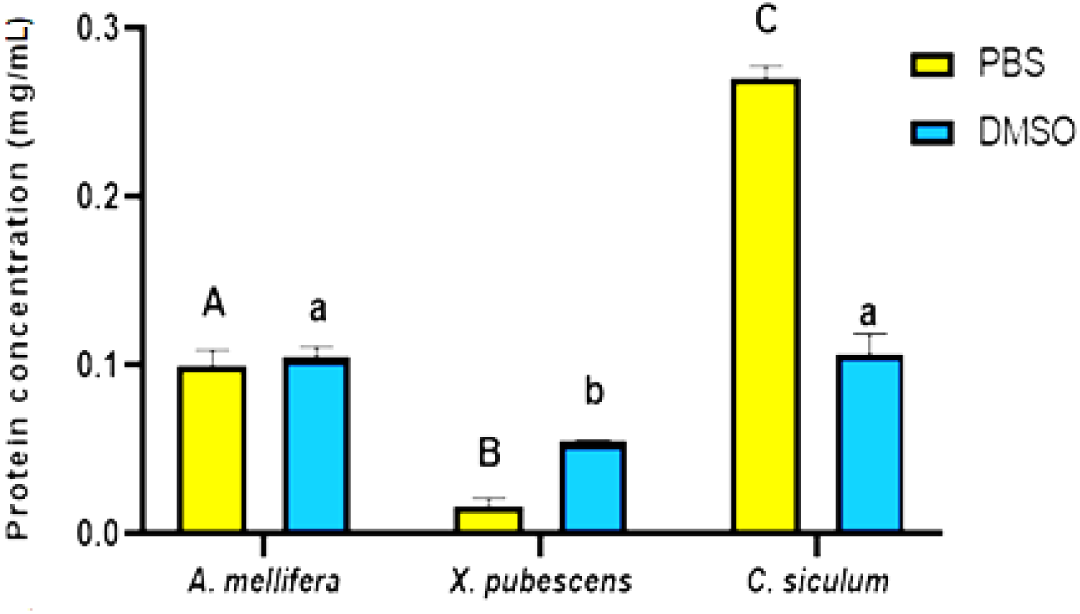
Protein concentration in mg/mL for hemolymph extracts of three bee species (*Apis mellifera, Xylocopa pubescens* and *Chalicodoma siculum*) in PBS or DMSO. Bars and error bars show the mean ± SE of triplicate results. Upper-case letters represent indicate significant differences between PBS extracts and lower-case letters indicate significant differences between DMSO extracts (Tukey’s HSD test: P ≤ 0.05).

## 2. Materials and Methods

### Bees

We sampled bees from 11 species (see details in Table 1) with a sweep net [19] between October 2020 (*A. mellifera*) and February 2021 (*X. pubescens* and *C. siculum*) in cultivated locations planted with faba beans (*Vicia faba*) in the Ismailia Governorate area (Egypt). We collected all specimens from each species in the same location on the same day. We then stored the bees at -20 °C until extraction and analysis of the hemolymph [20]. Since we only obtained sufficient amounts of hemolymph for three large-bodied species (*A. mellifera*, *X. pubescens*, *C. siculum*), we focused all our further analyses on these three species (Table 1).

**Table 1.** Details about bee sampling and hemolymph extraction.

| Bee species | Body size | Number of specimens | Weight of lyophilized hemolymph (grams) | Amount of hemolymph (microliter) | Location |
| --- | --- | --- | --- | --- | --- |
| *Apis mellifera* | Medium | 81 | 0.15 | ~800 | Ismailia |
| *Xylocopa pubescens* | Large | 20 | 0.34 | ~1700 | Ismailia |
| *Chalicodoma siculum* | Medium | 29 | 0.106 | ~590 | Ismailia |
| *Andrena savignyi* | Medium | 50 | <0.01 | < 50 | Ismailia |
| *Osmia latreillei* | Medium | <10 | <0.001 | < 50 | Ismailia |
| *Colletes lacunatus* | Medium | 16 | 0.039 | < 50 | Ismailia |
| *Amegilla quadrifasciata* | Medium | 60 | 0.0029 | < 50 | Ismailia |
| *Megachile flavipes* | Medium | 19 | <0.001 | < 50 | Ismailia |
| *Thyreus hyalinatus* | Large | 6 | <0.001 | < 50 | Ismailia |

### Hemolymph extraction and preparation

We extracted hemolymph from the specimen by using sterile insulin syringes to puncture the body close to the membrane of the coxa and applying a slight pressure on the abdominal region. We pooled the hemolymph samples from the same bee species and kept them in sterile Eppendorf tubes at -20 °C until lyophilization by using freeze drying [20]. We then dissolved the weighed lyophilized hemolymph samples in two different but complementary solvents (a hydrophilic solvent: phosphate buffer saline – PBS [21]; and a hydrophobic solvent: dimethyl sulfoxide – DMSO [22]) in order to extract a maximum of molecules as the polarity of the solvent influences the protein structure, solubility and stability [23]. For each test both PBS and DMSO were used as negative controls.

### Analysis of the protein content

#### Protein concentration

We measured the protein concentration (mg/mL) of the hemolymph extracts following Desjardins [24] using a Thermo Scientific™ (Waltham, MA, USA) Nano Drop™. One Micro volume UV-Vis Spectrophotometer with Bovine serum albumin (BSA) as standard in each sample (2 μL, 1 mg/mL). We analyzed three samples per bee species.

#### SDS-polyacrylamide gel electrophoresis

We used gel electrophoresis to separate protein bands of the hemolymph extracts based on their molecular weight using sodium dodecyl sulphate polyacrylamide gel electrophoresis (SDS-PAGE) following Laemmli [25] with modifications. We mixed equal volumes of hemolymph samples with solubilizing buffer (62.5 mM Tris-HCl (pH 6.8), 20% glycerol, 25(w/v) SDS, 0.5% 2-mercaptoethanol and 0.01% bromophenol blue). After heating for 4 min at 95°C, we inserted the samples into the wells (15 μL sample with 100 μg/mL protein concentration per well) of the separating gel (12% PAGE gel). We then ran an electrophoresis at constant 35 mA for 2 h using Consort N.V. (Belgium) mini vertical electrophoresis system with running buffer. We prepared the staining gel using 0.1% Coomassie Brilliant Blue (R-250) for the visualization of the protein bands.

#### High performance liquid chromatography

In complement to the electrophoresis, we performed an HPLC and analyzed the chromatograms by investigating the protein peaks and their separation based on their retention time. We obtained equal concentration (5 mg/mL) of hemolymph extracts in each of the two solvents (PBS or DMSO). We analysed the extracts (70 µL) using a YL9100 HPLC System with Stationary phase C18 column (Promosil C18 Column 5 μm, 150 mm×4.6mm) with acetonitrile (ACN) gradients of (10% - 100%) acetonitrile in water mobile phase for 50 min at flow rate = 1 mL/min. The chromatogram was detected using a UV detector at wavelength 280 nm following Basseri [26] with some modifications.

### Analysis of biological activities

#### Anticancer activity

We measured antitumor activities of hemolymph extracts using a 3-[4,5-methylthiazol-2-yl]-2,5-diphenyl-tetrazolium bromide (MTT) assay [27]. We seeded the cells from human liver cancer (HepG2) and human cervical cancer (HeLa) in 96-well plate for 24 h (5x103cells/well). After incubation, we treated the cells with 100 μL of the hemolymph extracts at serial concentrations in PBS and DMSO solvents (31.25, 62.5, 125, 250, 500 and 1000 μg/mL) and incubated them at 37 °C in 5% CO_2_ atmosphere for 48 h. We then washed the cells using PBS. We added fresh medium with MTT dye and incubated at 37°C for 4 h. We then added DMSO for the solubilisation of the formazan crystals in the viable cells [28]. We used a Bio-Tek ELISA micro plate reader to measure the absorbance at 540 nm. The experiment was performed in triplicates. We calculated the percentage of the cell viability (%) as follows:

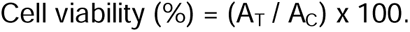

Where A_T_ was the absorbance of treated cells with extracts, and A_C_ was the absorbance of the control cells (untreated cells). The IC_50_ values (i.e. concentration of extracts that cause inhibition in the growth of 50% of the cells) were calculated for each sample using a dose-response curve with dose concentration (X-axis) and cell cytotoxicity percentage (Y-axis).

#### Antibacterial activity

We analyzed the antimicrobial activity of the hemolymph using the agar well diffusion method of Magaldi [29] against four bacterial strains (*Bacillus subtilis, Staphylococcus aureus, Escherichia coli* and *Salmonella typhimurium*). We inoculated the agar plate surface by distributing bacterial suspension. We then made a 6.0 mm hole aseptically using a sterilized tip and added 100 μL of each extract (10 mg/mL) or control into the well. We used PBS and DMSO as negative controls and Gentamycin as a positive control. After the incubation, we expressed the in vitro antimicrobial activity as inhibition zones in millimetres (mm) [30,31].

#### Scavenging ability

We measured the scavenging ability of hemolymph extracts using 1,1-Diphenyl-2-picrylhydrazyl (DPPH) free radical [32]. We incubated a mixture of 100μl DPPH methanolic solution (0.004% in 95% methanol) and 300μL of each hemolymph extracts at concentration 1 mg/mL and standard at 25 °C in the dark for about 30-60 min. We used ascorbic acid (Vitamin C) as positive control, and measured the colour changes using a spectrophotometer at 515 nm. We calculated the DPPH scavenging activity of the samples using the following equation [33]:

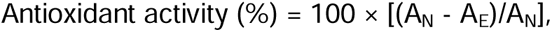

Where A_N_ was the absorbance of the negative control, and A_E_ was the absorbance of the sample or of the standard.

#### Hemolytic activity assay

We assessed the hemolysis activities of the bee hemolymph extracts against human erythrocytes by applying procedures of Malagoli [34]. We added blood samples to test tubes containing blood anticoagulant (EDTA) and centrifuged the tubes for 5 min at 10,000 rpm. We then suspended red blood cells in sterile PBS and incubated them with 100 μL volume from series of various concentrations of the tested extracts (range: 156.25 - 5000µg/mL). After incubation of the tubes at room temperature for one hour, we centrifuged the tubes for 5 min at 1x10^3^ rpm and measured the absorbance of the supernatant at 570 nm. We used Triton 10% as positive control while PBS and DMSO 10% as negative controls. We ran triplicate analysis for each bee species. We then calculated the hemolysis percentage for each extract as follows:

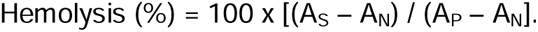

Where A_S_ was samples absorbance, A_N_ was the negative control absorbance, and A_P_ was the positive control absorbance.

### Statistical analysis

We analyzed the data in SPSS 22.0. We used Student’s t-tests to compare protein concentrations and IC_50_ of the hemolymph extracts in PBS and DMSO. We compared parameters of bee hemolymph (i.e protein concentrations, IC_50_s, antibacterial and antioxidant activities in either PBS or DMSO) across species using one-way ANOVAs followed by Tukey’s HSD post-hoc tests. We used two-way ANOVAs to compare parameters of bee hemolymph (i.e. protein concentrations, IC_50_s, antibacterial and antioxidant activities) in both solvents among the three bee species. We considered significant differences between samples when the p-value was lower than 0.05. All means are reported with their standard error (mean ± SE).

## 3. Results

### Analysis of protein content

#### Protein concentration

Protein concentration of hemolymph varied across species and solvents (Two-ways ANOVA, species x solvent: F (2, 12) = 111.97, P<0.001; Figure 1). Highest protein concentrations were recorded for *C. siculum* in PBS (0.27 ±0.01 mg/mL, n=3) and DMSO (0.107 ±0.01 mg/mL, n=3). Lowest protein concentrations were recorded for *X. pubescens* in PBS (0.02 ±0.005 mg/mL, n=3) and DMSO (0.06 ±0.001 mg/mL, n=3). Hemolymph of *A. mellifera* had values falling in between for PBS and similar values as *C. siculum* for DMSO.

#### SDS-polyacrylamide gel electrophoresis

Protein bands in electrophoresis gels of hemolymph extracted in PBS and DMSO had a molecular weight ranging from ∼5 to 250 kDa (Figure 2).

**Figure 2.**
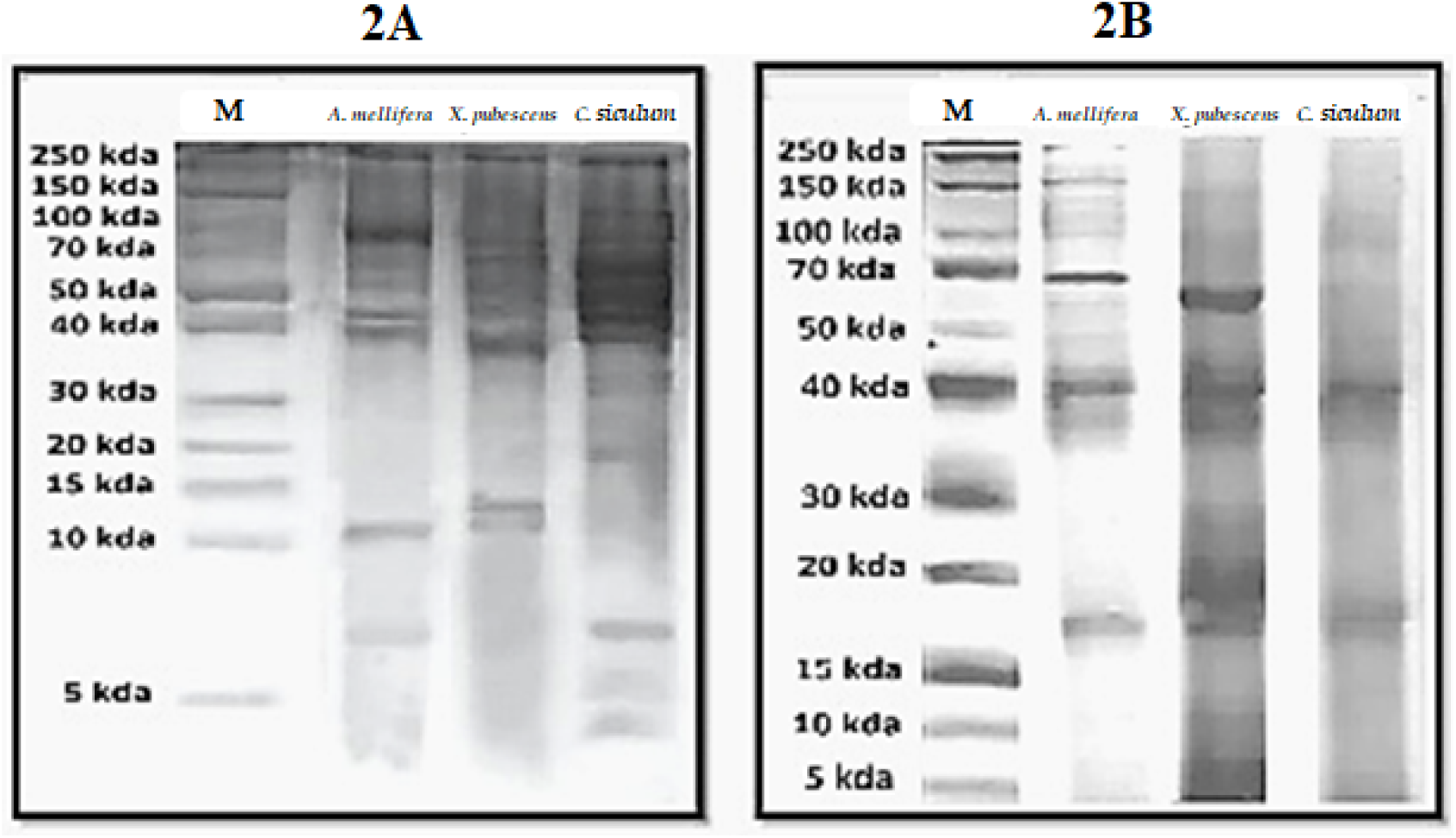
Pictures of SDS-PAGE gel for hemolymph of the three bee species (*Apis mellifera, Xylocopa pubescens* and *Chalicodoma siculum)* extracted in (A) PBS or (B) DMSO. M is the marker.

The PBS dissolved extracts were characterized by five bands common to the three bee species with molecular weights of ∼40, ∼60, ∼70, ∼110 and ∼250 kDa (Figure 2A). Other bands however were only observed in some species. A band with molecular weight of ∼7 kDa was only recorded in *A. mellifera* and *C. siculum*. Another band with molecular weight of ∼10 kDa was only recorded in *A. mellifera* and *X. pubescens*. Five bands with molecular weights of ∼4, ∼5, ∼17, ∼37 and ∼100 kDa were exclusively recorded in *C. siculum*.

The DMSO dissolved extracts were characterized by five bands common to the three bee species with molecular weights of ∼5, ∼17, ∼40, ∼60 and ∼110 kDa (Figure 2B). A band with molecular weight of ∼150 was recorded in *A. mellifera* and *X. pubescens*. Two bands with molecular weights of ∼10 and ∼25 kDa were only recorded in *X. pubescens*.

#### RP-HPLC analysis

By analyzing the chromatogram of the PBS dissolved hemolymph extracts we found the presence of 6 peaks common to the three bee species at retention times of 7.5, 11.1, 14.4, 15.4, 22.9 and 28.3 min (Table 2). In DMSO dissolved extracts, we detected 10 peaks common to the three bee species at retention times of 9.5, 23.3, 24.6, 25.5, 26.8, 28.3, 32.6, 33.2, 40.3 and 44.8 min (Table 2).

**Table 2.** HPLC most common chromatogram peaks profiles of the hemolymph of three bee species (*Apis mellifera, Xylocopa pubescens* and *Chalicodoma siculum*) extracted in PBS or DMSO. N.A.: not applicable.

| Retention time<br>(minutes) | Peak area ( $\times 10^2$ mV.s) | | | | | |
| --- | --- | --- | --- | --- | --- | --- |
|  | *A. mellifera* |  | *X. pubescens* |  | *C. siculum* |  |
|  | PBS | DMSO | PBS | DMSO | PBS | DMSO |
| 7.5 | 4.6 | N.A | 0.7 | N.A | 177.4 | N.A |
| 9.5 | N.A | 327 | N.A | 7.7 | N.A | 9.7 |
| 11.1 | 2.3 | N.A | 0.4 | N.A | 25.2 | N.A |
| 14.4 | 5.9 | N.A | 0.3 | N.A | 10.3 | N.A |
| 15.4 | 5.5 | N.A | 0.4 | N.A | 2.2 | N.A |
| 22.9 | 13.9 | N.A | 1.2 | N.A | 3.5 | N.A |
| 23.3 | N.A | 3.6 | N.A | 0.1 | 3.6 | 3.4 |
| 24.6 | N.A | 3.4 | N.A | 0.6 | N.A | 1.4 |
| 25.5 | 9.1 | 3.6 | 1.5 | 0.1 | N.A | 3.8 |
| 26.8 | N.A | 4.0 | 1.1 | 0.3 | 2.8 | 0.6 |
| 28.3 | 5.3 | 1.5 | 1.5 | 0.4 | 1.2 | 0.7 |
| 32.6 | N.A | 1.4 | N.A | 2.2 | N.A | 1.0 |
| 33.2 | N.A | 0.3 | N.A | 1.3 | N.A | 0.8 |
| 40.3 | N.A | 0.1 | N.A | 0.1 | N.A | 0.1 |
| 44.8 | N.A | 1.0 | N.A | 1.2 | N.A | 0.9 |

### Analysis of biological activities

#### Anticancer activity

We evaluated the antiproliferative actions of the hemolymph extracts against the viability of hepatic and cervical carcinoma cells. All hemolymph extracts resulted in inhibition of the cell viability against the tested cancer cell lines in a dose-dependent manner after 48 h of incubation, irrespective of the solvent used (Figure 3). However, the DMSO dissolved extracts showed higher overall cytotoxic activity than the PBS dissolved extracts for HepG2 and for HeLa (Two-ways ANOVA, HepG2: F (2, 12) = 26.667, P<0.001; HeLa: F (2, 12) = 117.111, P<0.001) (Figure 4).

**Figure 3.**
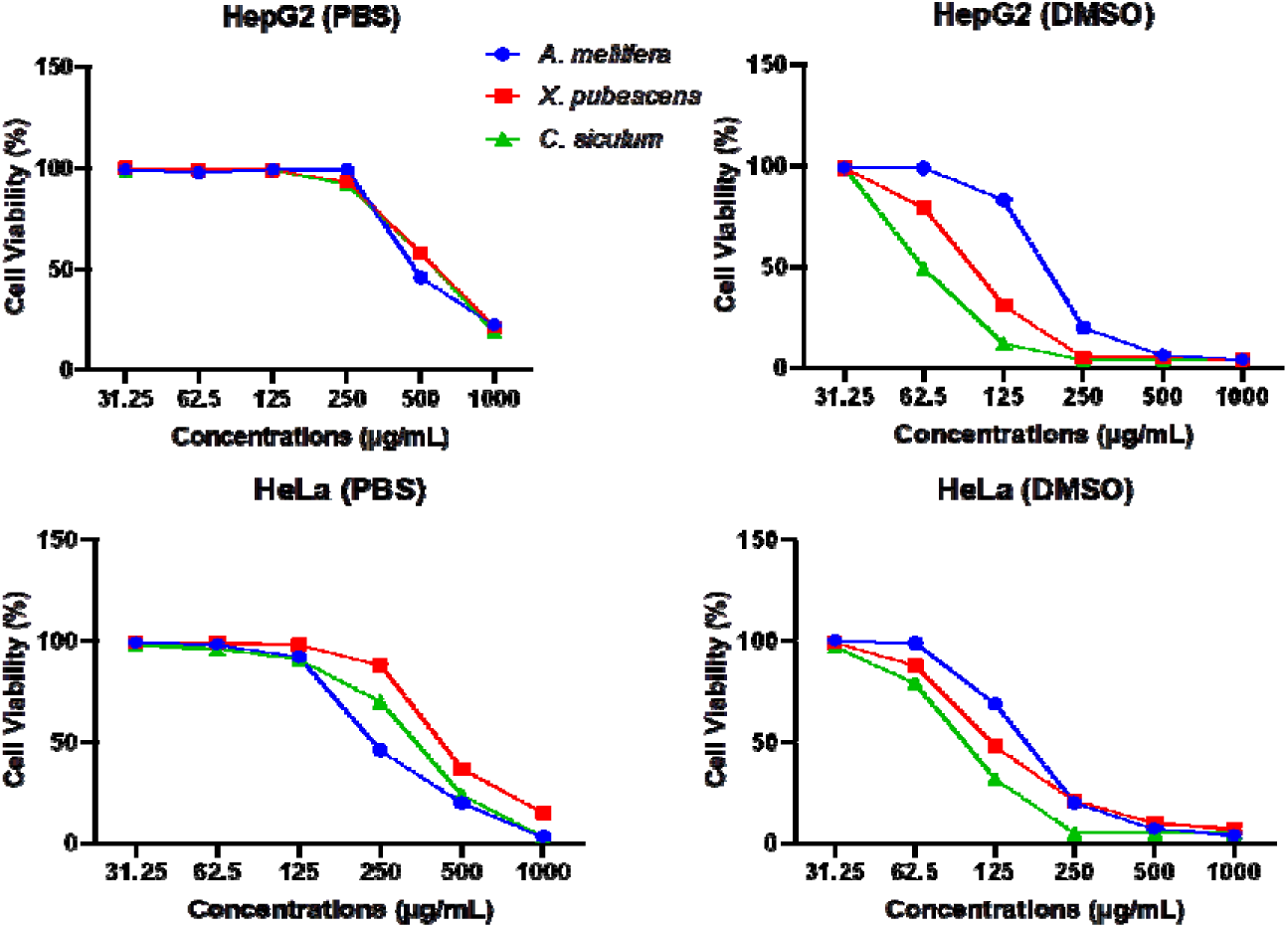
Effects of hemolymph extracts of three bee species (*Apis mellifera, Xylocopa pubescens* and *Chalicodoma siculum*) on cell proliferation of human cancer (HepG2 and HeLa) cell lines at different concentrations in PBS or DMSO.

**Figure 4.**
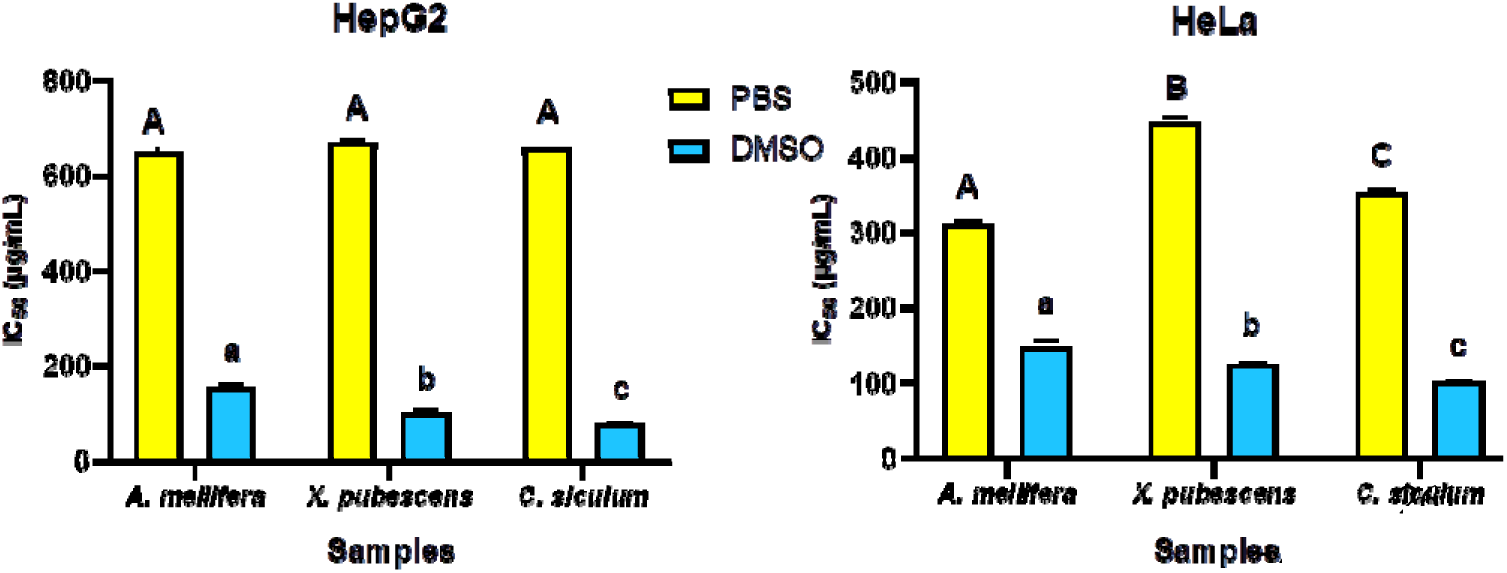
IC_50_ in µgram/mL of hemolymph extracts of three bee species (*Apis mellifera, Xylocopa pubescens* and *Chalicodoma siculum*) against HepG2 and HeLa cell lines, using PBS or DMSO as solvents. Bars and error bars represent the mean values ±SE of triplicate results. Upper-case letters indicate significant difference between PBS extracts and lower-case letters indicate significant differences between DMSO extracts (Tukey’s HSD test: P ≤ 0.05).

Among the DMSO dissolved extracts, *C. siculum* hemolymph had the highest cytotoxic effects against HepG2 and HeLa cell lines (IC_50_ HepG2 = 77.35 µg/mL, IC_50_ HeLa = 101.2 µg/mL). The lowest cytotoxic effects were recorded for *A. mellifera* hemolymph (IC_50_ HepG2 = 153.1 µg/mL, IC_50_ HeLa = 148.46 µg/mL) (Figure 4). Among the PBS dissolved extracts, *A. mellifera* hemolymph had the highest cytotoxic effects (IC_50_ HepG2 = 649.4 µg/mL, IC_50_ HeLa = 312.54 µg/mL) and *X. pubescens* had the lowest (IC_50_ HepG2 = 669.2 µg/mL, IC_50_ HeLa = 447.2 µg/mL) (Figure 4).

#### Antimicrobial activity

We observed the antibacterial activity of hemolymph extracts against Gram-positive and Gram-negative bacteria (Table 3, see example in Figure 5). The results showed comparable antibacterial activities of hemolymph extracts among the tested bee species and solvent; *Bacillus subtilis* (ANOVA, F (2, 12) = 0.800, P=0.472), *Staphylococcus aureus* (ANOVA, F (2, 12) = 0.042, P=0.960), *Escherichia coli* (ANOVA, F (2, 12) = 2.054, P=0.171) and *Salmonella typhimurium* (ANOVA, F (2, 12) = 2.771, P=0.102).

**Figure 5.**
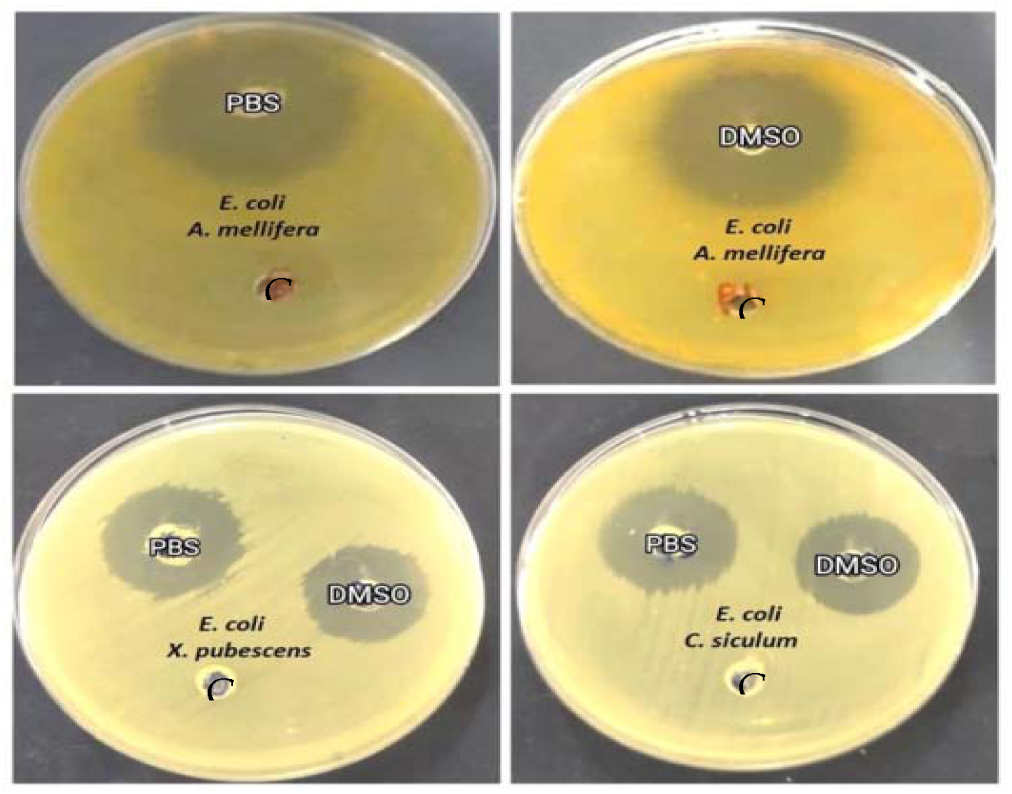
Antimicrobial activity of hemolymph from three bee species (*Apis mellifera, Xylocopa pubescens* and *Chalicodoma siculum*) extracted in PBS or DMSO against *E. coli* bacteria and by using negative control.

**Table 3.**
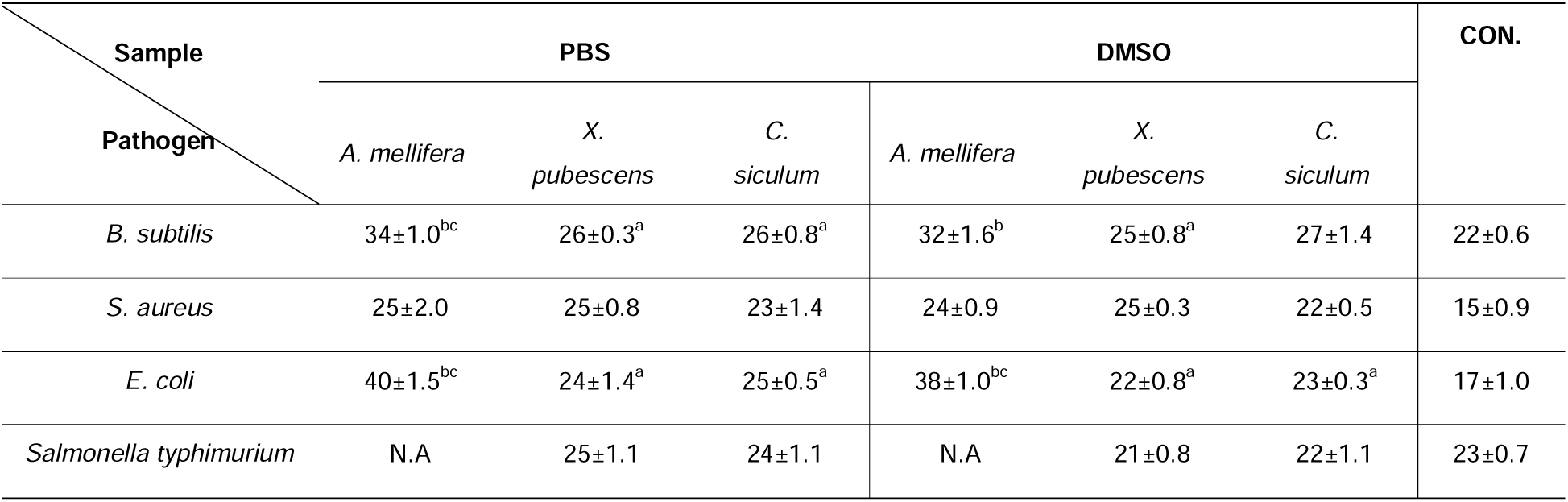
Inhibition zone (mm) of hemolymph from three bee species (*Apis mellifera, Xylocopa pubescens* and *Chalicodoma siculum*) extracted in PBS or DMSO against various types of bacteria (*Bacillus subtilis, Staphylococcus aureus, Escherichia coli* and *Salmonella typhimurium*). CON: Gentamycin as positive control, N.A: not applicable, ±S.E.: standard error values obtained from triplicate measurements. Values with different superscript letters in a column are significantly different (Tukey’s HSD test: P ≤ 0.05), a versus *A. mellifera*, b versus *X. pubescens* and c versus *C. siculum*.

#### DPPH radical scavenging assay

The hemolymph extracts of *A. mellifera, X. pubescens* and *C. siculum* possessed effective scavenging actions in both PBS and DMSO (Figure 6). Overall, the hemolymphs of the different species showed different antioxidant activities (two-ways ANOVA, species: F (2, 12) = 5.06, P=0.025) and these values were higher in PBS dissolved extracts (two-ways ANOVA, solvents: F (1, 12) = 10.729, P=0.007). However, for a given species the antioxidant activity in both solvents was comparable (two-ways ANOVA, species x solvent: F (2, 12) = 2.451, P=0.128).

**Figure 6.**
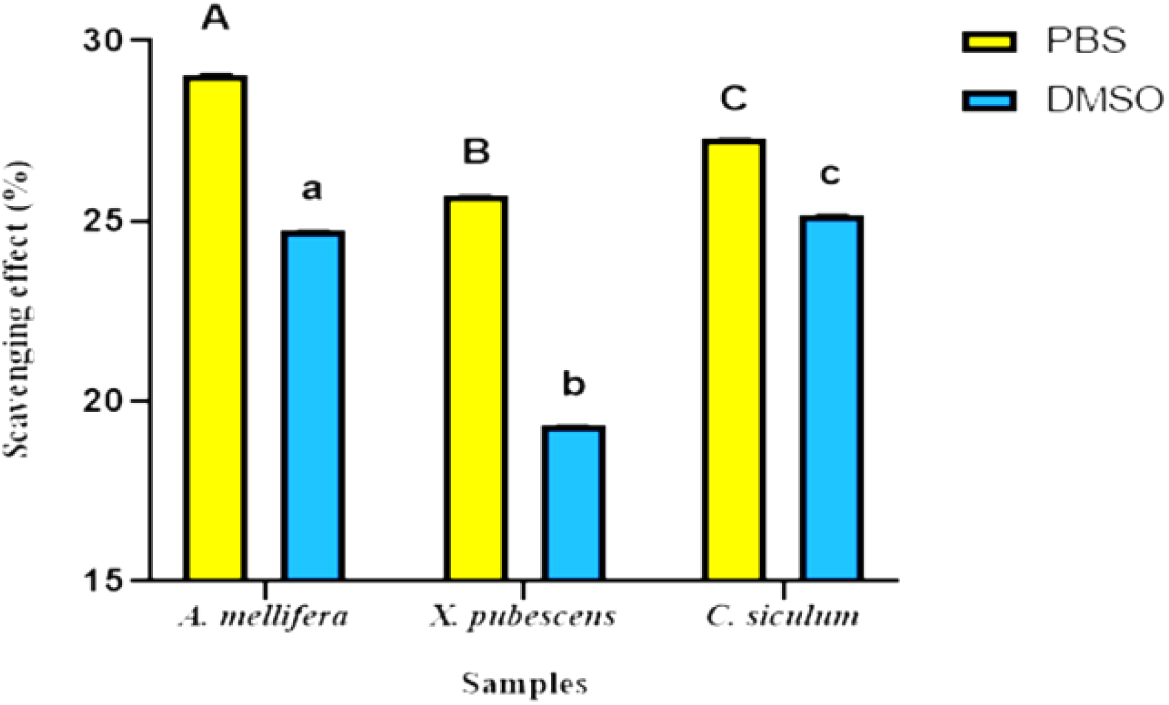
DPPH radical scavenging activity of hemolymph extracts of three bee species (*Apis mellifera, Xylocopa pubescens* and *Chalicodoma siculum*) extracted in either solvent PBS or DMSO. Bars and error bars denote the mean values ± S.E. of triplicate results. Upper-case letters indicate significant differences between PBS extracts and lower-case letters indicate significant differences between DMSO extracts (Tukey’s HSD test: P ≤ 0.05).

The highest antioxidant activity was reported for *A. mellifera* PBS extracts (29.1%) and *C. siculum* DMSO extracts (25.2%), while *X. pubescens* reported the lowest antioxidant activity (25.7%) in PBS and (19.3%) in DMSO (Figure 6).

#### Hemolytic activity

Hemolytic properties of the bee hemolymphs were tested against human erythrocytes. For the three bee species, hemolymph extracts possessed either no or low hemolytic activity against human erythrocytes in reference to negative (PBS and DMSO 10%) and positive controls (Triton10%) (Figure 7).

**Figure 7.**
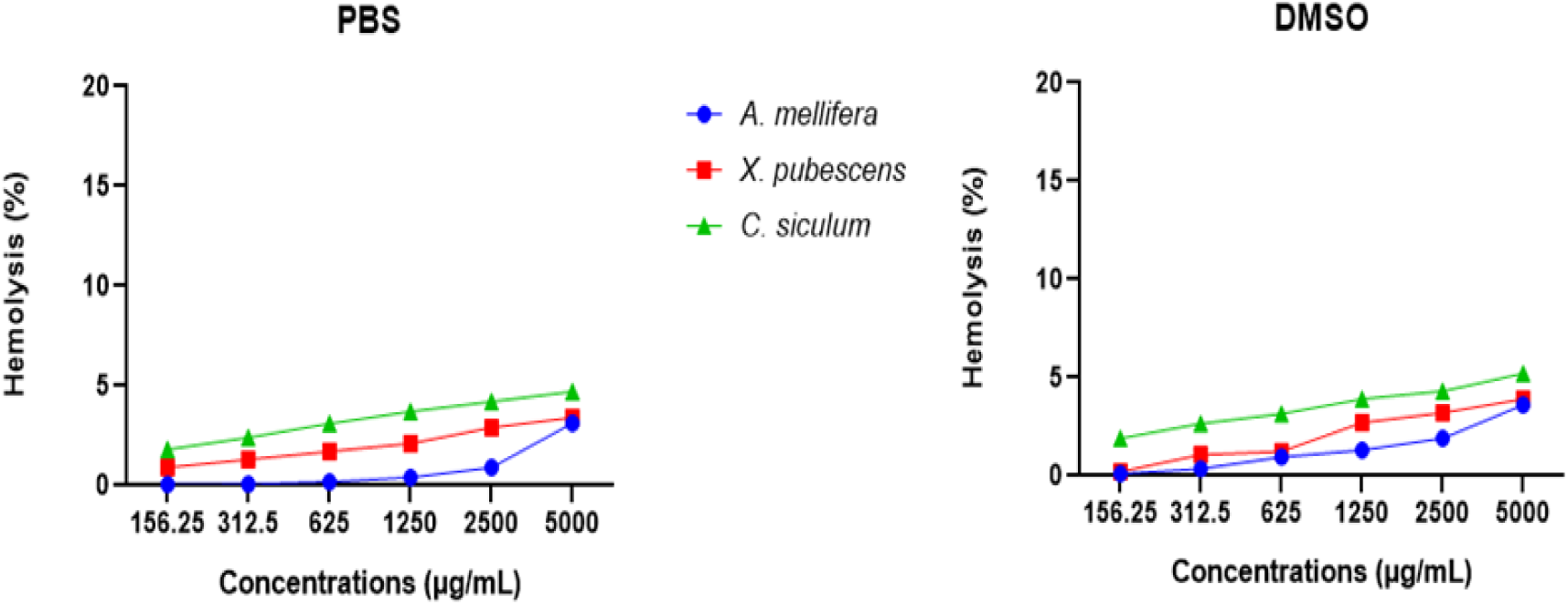
Hemolytic activity of hemolymph extracts from three bee species (*Apis mellifera, Xylocopa pubescens* and *Chalicodoma siculum*) at different concentrations against human erythrocytes. Hemolymph samples were extracted in PBS or DMSO.

## 4. Discussion

We aimed to evaluate the interspecific variability in hemolymph protein content and biological activities from bees collected in the same locality in Northern Egypt. We found considerable variation among the three bee species for which we managed to extract enough hemolymph after our sampling campaign (see summary results in Table 4).

**Table 4.** Summary of analysis of hemolymph of three bee species (*Apis mellifera, Xylocopa pubescens* and *Chalicodoma siculum*) extracted in two solvents (PBS and DMSO). (+): high, (+/-): medium and (-): low.

|  | *A. mellifera* |  | *X. pubescens* |  | *C. siculum* |  |
| --- | --- | --- | --- | --- | --- | --- |
|  | PBS | DMSO | PBS | DMSO | PBS | DMSO |
| Protein concentration | (+/-) | (+/-) | (-) | (-) | (+) | (+/-) |
| Anticancer activity | (+) | (-) | (-) | (+/-) | (+/-) | (+) |
| Antimicrobial activity | (+) | (+) | (+/-) | (+/-) | (+/-) | (+/-) |
| Antioxidant activity | (+) | (+/-) | (-) | (-) | (+/-) | (+) |
| Hemolytic activity | (-) | (-) | (-) | (-) | (-) | (-) |

Overall*, X. pubescens* hemolymph possessed the lowest protein concentrations, *A. mellifera* hemolymph had intermediate values, and *C. siculum* hemolymph showed highest protein concentrations irrespective of the solvent used for extraction. Previous studies reported intraspecific variations in hemolymph protein concentration primarily correlated to the diet of bees, even among bees from the same worker caste [6,35–37]. Since all our samples came from the same study site and were collected on the same plants, this suggests *A. mellifera, X. pubescens* and *C. siculum* had different diets. Bee diet varies based on various parameters [38] including the physiological states of the bees themselves and the developmental stages of larvae they need to feed [39–41]. Another source of variation is the impact of presence of phoretic mites on Xylocopa species that feeds on pollen paste [42], the main source of proteins in the hemolymph of bees [43].

The lowest protein concentrations in *X. pubescens* were associated with the weakest antioxidant and antibacterial activities against both *B. subtilis* and *E. coli*. This was also associated with the weakest anti-proliferative activities among the PBS dissolved hemolymph extracts against both HepG2 and HeLa cell lines. By contrast, the highest protein concentrations of *C. siculum* in both solvents were associated with the strongest anti-proliferative activities among DMSO dissolved hemolymph extracts against both tested cancer cell lines. This is consistent with the fact that high protein concentration in the bee hemolymph is associated to high resistance to pathogens, increased life span, and improved immunity [11–13]. Therefore protein concentration in hemolymph is a good indicator of bee health.

SDS-PAGE gel analysis revealed proteomic profiling of the hemolymph extracts of the three bee species in PBS or DMSO solvents. Protein bands common to the three species, with molecular weight (4-5 kDa) in DMSO dissolved hemolymph were similar to cercopin, a family of small proteins have been isolated from different insect hemolymph which possesses anti-bacterial properties against both Gram positive and Gram negative bacteria [44,45]. Extractions in DMSO also revealed another protein bands common to the three species that were detected at molecular weight (lil17 kDa) similar to a group of lectins [46]. Lectin is a defense protein that can participate in many biological activities such as antimicrobial, antioxidant and anticancer in arthropods. It can be also considered as natural anticancer agent and contribute in immune actions [47–53]. CLIPC9 similar bands were observed at (lil37 kDa) in both PBS and DMSO extracts consistent with their detection in the hemolymph of *Anopheles gambiae* [54]. CLIP proteases are serine proteases found in the hemolymph of all insects and participate in the innate immune responses. The protein bands with molecular weight (lil40 kDa) were also found in both PBS and DMSO extracts are in the same range of lipopolysaccharide recognition protein (LRP). LRP was purified from the plasma of large beetle larvae, *Holotrichia diomphalia* with immune roles as it participates in agglutinating activities against *E. coli* and other bacteria [55]. Protein bands detected at molecular weight (lil60 kDa) in PBS or DMSO extracts are within the range of purified protein fractions extracted from cockroach hemolymph with potent antimicrobial activities [26]. Moreover, common detected protein bands at (lil70 kDa) were similar to hemocyanin, an oxygen transporter protein that possesses antioxidant, antiparasitic, antimicrobial, anticancer and other biotic activities. This protein is found in most arthropods [53,56,57].

The variety and likely dominance of the bioactive proteins such as cercopins, lectins, CLIP proteases, LRP and hemocyanin in PBS and DMSO hemolymph extracts may clarify the effective biological activities of hemolymph in the current study. Generally, DMSO dissolved hemolymph extracts possessed higher cytotoxic activities against HepG2 and HeLa cancer cell lines than the effect of PBS extracts. Our proteomic profiling analysis revealed the presence of variations according to the type of dissolved proteins in different solvents, in agreement with studies on the impact of solvent on the protein structure [21,58]. According to Kramer [59] and Nugraha [21], altered solvents affects the protein solubility thus may influences the hemolymph bio-activities. Hydrophobic proteins such as lipoprotein complexes were used as delivery vehicles for anticancer drugs [60].

The fact that none of the hemolymph extracts possessed lytic activities against erythrocytes agrees with results of previous study on the hemolytic activities of honey bee hemolymph [6]. These results may encourage the evaluation of bee hemolymph extracts as safe and selective therapeutic agents. Accordingly, bee hemolymph mixed with herbal extracts exhibited high anticancer efficiency with less or no hemolytic activities [4].

Honey bees have innate immune mechanisms including the physical barriers and both humoral and cellular actions for their defense against infections and pathogens that affects the bee immune system and thus affects the bee health and the social behavior of these insects develops their social immunity which reduces the stress of the individual immune response of the bees [61]. Further studies are required on the immune responses of other different bee species for explaining the variations in their proteomic content and biological activities

## 5. Conclusions

Our study highlights significant interspecific variability in the protein concentration and biological activities of the hemolymph of three common Mediterranean bee species, likely related to variation in their life history traits. Broader scale comparative studies, using similar procedures as ours, across different geographical areas are now needed to investigate variations among bee species and how they are related to their ecology and evolution.

## Author Contributions

**MT, IB and MS:** Conceptualization and Supervision**. SE, MT, IB and MS:** Methodology, Software, validation, formal analysis, investigation, resources and data curation**. SE, MT, IB, MS and ML:** Writing—Original Draft Preparation, Writing—Review & Editing, visualization. All authors have read and agreed to this version of the manuscript.

## Data Availability Statement

Data available on request due to restrictions, e.g., privacy or ethical.

## Acknowledgments

We would like to thank Soliman Kamel and Saied Aboud for their contribution in the bee sample collection.

